# Explorations of the polygenic genetic architecture of flowering time in the worldwide *Arabidopsis thaliana* population

**DOI:** 10.1101/206706

**Authors:** Yanjun Zan, Örjan Carlborg

## Abstract

As a locally adapted complex trait, flowering time in *Arabidopsis thaliana* has attracted much attention in genetics. Most studies have, however, focused on contributions by individual loci rather than the joint contributions by the large number of loci in the genetic architecture of flowering time to local and global adaptation. In an earlier study, we reported 46 loci associated with flowering time variation during growth at 10°C, 16°C or both in the 1,001-genomes collection of *Arabidopsis thaliana* accessions. Here, we explore how these loci together contribute to differences among genetically defined, and geographically divided, subpopulations across the native range of this species. Our approach was to define flowering time as a trait, and the measurements at 10 and 16 °C as two independent measurements of it. This facilitated explorations of the dynamics in the genetic architecture –which loci contribute and their effects– of flowering time across growth temperatures and their potential roles in local and global adaptation. The overall flowering time differences between populations could be explained by subtle changes in allele-frequencies and gradual changes in phenotype due to globally present alleles. More extreme local adaptations were on several occasions due to contributions by regional alleles with relatively large effects. About 2/3 of the 48 evaluated flowering time loci had similar effects on flowering time at 10°C and 16°C, while the remaining 1/3 had different effects in the two temperatures, suggesting an important contribution of gene by temperature interactions to this trait. There are also indications that co-evolution of functionally connected alleles in local populations has been important for local adaptation. Overall, this study provides deeper insights to the polygenic genetic basis of flowering time variation in *Arabidopsis thaliana* across a wide range of ecological habitats.

**Author Summary:** Many genes can affect flowering time in *Arabidopsis thaliana*, but their contribution to natural flowering time variation in the worldwide population is largely unknown. We explored how 48 loci associated with flowering time, measured at 10°C and 16°C, or their difference, for the same wild collected 1,001-genomes *Arabidopsis thaliana* accessions together contribute to differences among the genetically defined and geographically divided subpopulations from the native species range. The overall flowering time differences among these subpopulations could be explained by the joint small effects of globally present alleles, suggesting an important contribution by polygenic adaptation for this trait. Most alleles with large effects on flowering were present only in some populations, facilitating more extreme local adaptations. Long-range LD was observed between genes in several biological pathways, indicating possible local adaptation via co-evolution of functionally connected polymorphisms. The genetic architecture of flowering time was also found to depend on the growth temperature. Most flowering time loci had similar effects on flowering time measured at 10°C and 16°C, but the effects of about 1/3 of them had effects that varied with temperature. Overall, new insights are provided to how the polygenic architecture of flowering time has facilitated its colonisation of a wide range of ecological habitats.

## Introduction

Understanding the genetic basis of adaptation is a central theme in evolutionary genetics [1, 2]. Adaptation in nature can rely on a few loci with large individual effects [1, 3], and such loci have been identified in natural populations of for example *Arabidopsis thaliana* [4–7] and the Darwin finch [8]. In other cases, populations adapt via selection on many loci with small effect [9], with examples reported for fitness traits in a range of species including mice, drosophila, human and maize[10–12]. In depth characterizations of the genetic mechanisms contributing to adaptation for polygenic quantitative traits is, however, a difficult [13] and despite some successes little is overall known. Adaptations for polygenic traits could be rapid due to selection on available standing variation [14–16] and new adaptive optima could be reached via gradual allele frequency changes across multiple loci [17, 18]. The individual effects of the contributing loci might be too low for detection in genome wide scans based on stringent multiple testing thresholds, but they might despite this be necessary for reaching fitness optima [19–21].

*Arabidopsis thaliana*, a widely used model species in plant biology, has colonised a broad range of ecological habitats around the world. One of the most well studied adaptive traits is flowering time due to its role in ecological adaptation and potential impact on agronomic production in related species. Molecular studies in the laboratory, primarily using the reference accession *Col-0*, have identified many genes with the potential to alter flowering time. Association studies in natural populations, however, rarely reveal more than a handful of the loci contributing to the genetic variation of this trait [22–25]. Hence, even though it is likely that flowering time adaptations across the native range of *A. thaliana* would involve many loci, the polygenic basis of these differences is still largely unexplored.

Previously, we developed and used a new statistical approach to dissect the genetic basis of polygenetic traits in natural populations. The properties of the method was illustrated by exploring the polygenic basis of flowering time in *Arabidopsis thaliana*, using public data from the 1,001 genomes project[26]. In total, 33/29 loci were there mapped for flowering times in 1,004 wild collected *Arabidopsis thaliana* accessions from the 1,001-genomes project grown at 10/16°C[26]. Here we explored how the genetic effects of these loci i) vary within and across loci and growth temperatures and ii) how these variations in allelic effects contribute to the flowering time variation between genetically defined subpopulations that have colonized the native range of the species [22].

Loci with globally segregating alleles made, in general, small individual contributions to flowering but together captured the overall pattern of flowering time differences between the subpopulations, suggesting a role of polygenic adaptation[14–18]. The alleles with more restricted geographic distributions had in several instances large effects and made important contributions to local adaptations of some subpopulations. The genetic effects of many loci varied across temperatures, suggesting that genotype by temperature interactions might be important for flowering time adaptation. The long-range LD between polymorphisms in genes of the same biological pathways suggests that co-evolution of different pathways has contributed to local adaptation in different geographic areas. Overall this reanalysis of public data provides new insights to the polygenetic basis of flowering time adaptation in *A. thaliana.*

## Results

### Flowering time differentiation in *A. thaliana* worldwide population

Flowering time variation of *Arabidopsis thaliana* accessions sampled from its native range is strongly associated with geographic variation in climate [27–29]. To study the genetics of the this geographical differentiation in greater detail, we reanalysed a publicly available dataset where flowering time was measured at 10°C (FT10) and 16°C (FT16) under greenhouse conditions on 1,004 whole-genome re-sequenced *Arabidopsis thaliana* accessions[22]. The genetic population structure and geographic sampling locations for the analysed accessions are illustrated in S1 Fig.

We approached flowering time in this population as a single complex trait measured at two temperatures to reveal the dynamics in its underlying genetic architecture. At these temperatures, flowering times displayed a continuous variation with earlier average flowering times (9.3 days) at 16 °C than at 10 °C (Figure 1A). The variations in flowering times at both growth temperatures were correlated with the location and climate at the sampling sites of the accessions, suggesting its potential role in global, and local, adaptation. For example, there was a significant latitudinal cline of flowering times (Fig 1B/S2A Fig for FT10/FT16; Pearson correlation = 0.37; P < 2.2 × 10^−16^; df = 963) and a significant correlation between flowering times and mean temperature at the sampling locations (Fig 1C/S2B Fig for FT10/FT16; Pearson correlation = −0.31; P < 2.2 × 10^−16^; df = 963).

**Fig 1.**
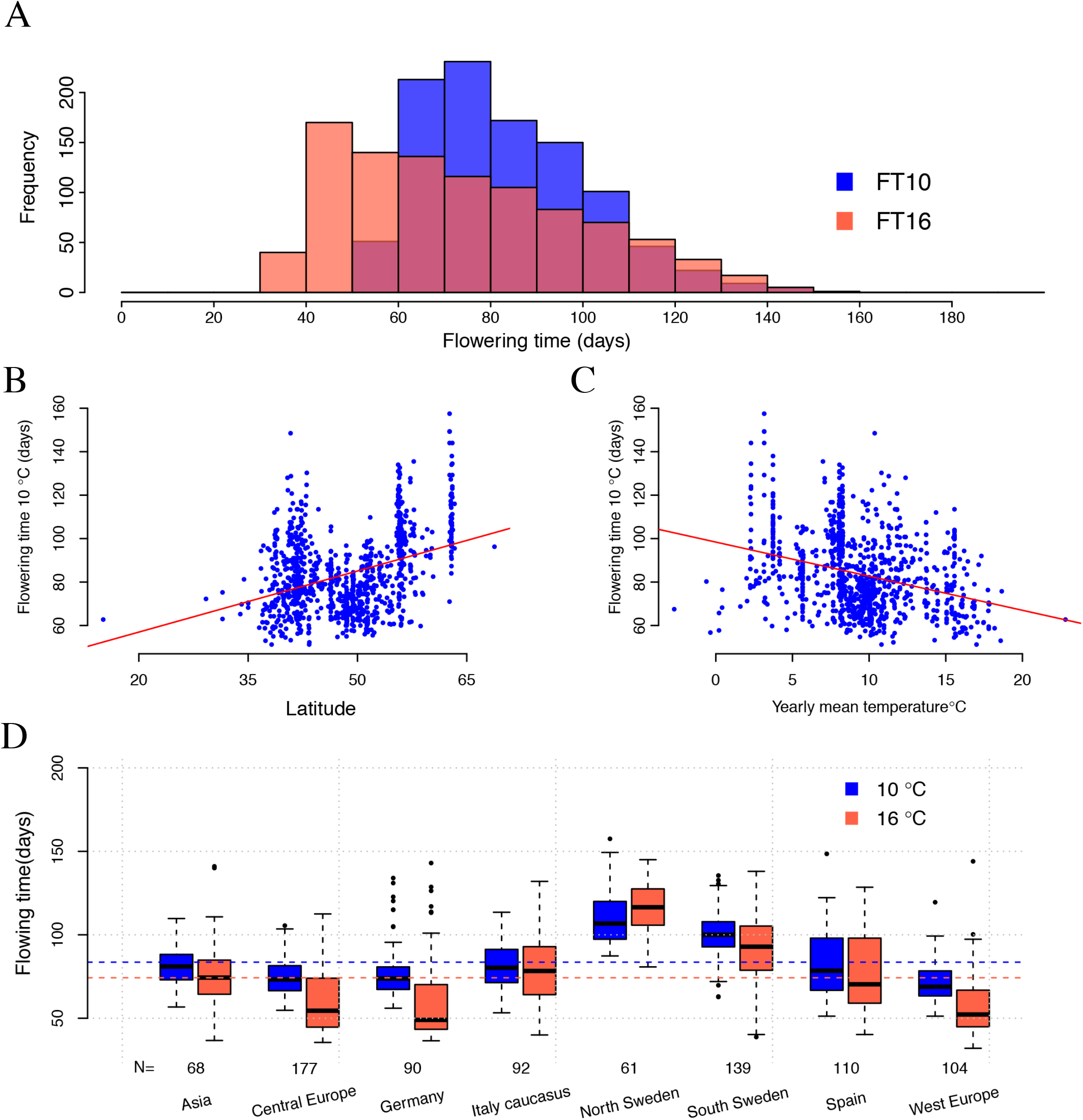
Flowering time measured at 10 and 16 °C, correlation with latitude/temperature, and the subpopulation differentiation. **(A)**. Histogram of flowering time measured at 10°C (blue) and 16 °C (tomato). Flowering time under controlled green house condition is significantly correlated both with latitude **(B)** and temperature **(C)** at the sampling sites, and vary amongst the earlier defined subpopulations **(D)**, of the 1,004 analysed accessions from the 1,001-genomes *A. thaliana* collection [22].

The phenotypic variance increased with temperature, being larger at 16°C than at 10°C (Fig 1A), while the proportion of additive variance estimated by fitting the IBS (Identity By State) kinship matrix were similar at both temperatures (82/83% at 10/16°C). Additive genetic variance is thus released as the temperature increases, which will be explored in more detail in the following section. Regardless of whether flowering times were measured at 10°C or 16°C, there were substantial phenotypic differences between the eight genetically defined and geographically separated subpopulations defined by *Alonso et al* [22] in their admixture analysis of this population (Fig 1D). Increasing the growth temperature leads to earlier flowering times in most of the sub-populations, with the North Swedish population being the only exception with an on average later (4.9 days) flowering time at 16°C.

### The genetic architecture of flowering time is highly polygenic in the worldwide *A.thaliana* population

By utilizing prior molecular knowledge of genes regulating flowering time variation in the reference accession *(Col-0)*, and results from an expression QTL analysis of those genes in this 1001-genomes collection of natural accessions, we earlier mapped 46 flowering time loci where 33/29 loci were associated with FT10/FT16 using a multi-locus association analysis[26]. Together these mapped loci explain 55/48% of the total genetic variance for FT10/FT16[26]. Here, rather than focusing on the results from the two temperatures separately, the complete genetic architecture of flowering time revealed across them was evaluated. Overall, 16 loci were mapped at both temperatures, and 2 pairs of loci were located within 20 kb from each other, suggesting that they contribute to flowering time regardless of temperature in a relatively stable manner. The remaining 30 loci were significantly associated with flowering times only at one temperature in the earlier analysis[26], and their possible involvement in genotype-by-temperature interactions are further explored here.

Two additional analyses were performed to explore the possible role of genotype-by-temperature interactions in this population. First a GWA mapping analysis to the flowering time difference at the two evaluated temperatures (FT10-FT16). Two genome-wide significant associations, one at *Chr1: 23,453,207* bp and one at *Chr4: 1,338,596* bp, were detected in this analysis after Bonferroni correction for ∼1.4 million SNPs (S3 Fig). Second, the polygenic contribution of the loci that were significant in one temperature, but not the other, to flowering times was evaluated. This by testing if loci associated with FT16, but not FT10, together with the two additional loci affecting FT10-FT16 jointly contributed a significant amount of genetic variation in FT10, and vice versa. Models with or without nonoverlapping loci were fitted, and significant increases in the adjusted R^2^ were detected at both temperatures (4/8% for FT10/FT16; Likelihood ratio test; P-value= 3.2 × 10^−14^/1.3 × 10^−33^; S4 Fig).

Linear models including the genotypes of the 48 mapped flowering time loci, but excluding their kinship, explained the majority of the additive genetic variance at both temperatures (79/75% for FT10/FT16). In the following sections, we will explore how these 48 loci together contribute to flowering time across the different genetically and geographically divided sub-populations in this population, to potentially gain deeper insights to how polygenic and genotype-by-temperature interactions might contribute to local and global adaptation in nature.

### The role of genotype-by-temperature interactions for flowering times in *A. thaliana*

It is known that genotype-by-temperature interactions are important for flowering times in *Arabidopsis thaliana* [30–32]. The genetic basis of this phenomenon is, however, largely unknown. Here, we studied this by evaluating the temperature dependence of the genetic effects of 48 mapped flowering time loci. As shown in Fig 1A, increased growth temperature leads to a release of additive genetic variance in flowering time. To identify the genetic basis of this, the genetic effects of the 48 mapped flowering time loci on FT10 and FT16 were estimated by fitting them jointly in a mixed model (Fig 2). The alleles at 42 of the 48 loci have effects in the same direction on FT10 and FT16, i.e they consistently increase or decrease flowering time regardless of temperature. The alleles at the remaining six loci have opposite effects in 10°C and 16°C, but as all these estimates in at least one temperature are close to zero (Fig 2A). This result is likely to at least in part be the outcome of the uncertainty in the estimation of the allelic effects. For ∼2/3 of the loci, the sizes of the genetic effects on flowering time are in good general agreement (Fig 2A), suggesting that they affect flowering time in similar way regardless of whether the plants are grown at 10°C or 16°C. The remaining ∼1/3 loci, however, have larger effects on flowering time when the plants are grown at 16°C than when they are grown at 10°C (Fig 2A). This is consistent with the observation of an increased additive genetic variance at 16°C, but at the same time unexpected as an increase in temperature generally leads to an overall decrease in flowering time. To evaluate whether these putative genotype-by-temperature were statistically significant, we compared the fits of two linear models for every locus: one including all 48 loci and an indicator variable for the two growth temperatures as fixed effects, and another also including an interaction term between growth temperature and each tested locus. In total, 18 loci displayed significant genotype-by-temperature interactions after correction for multiple testing (highlighted in Fig 2A; names of loci & P-values from the corresponding Likelihood Ratio Test in Table S1).

**Fig 2.**
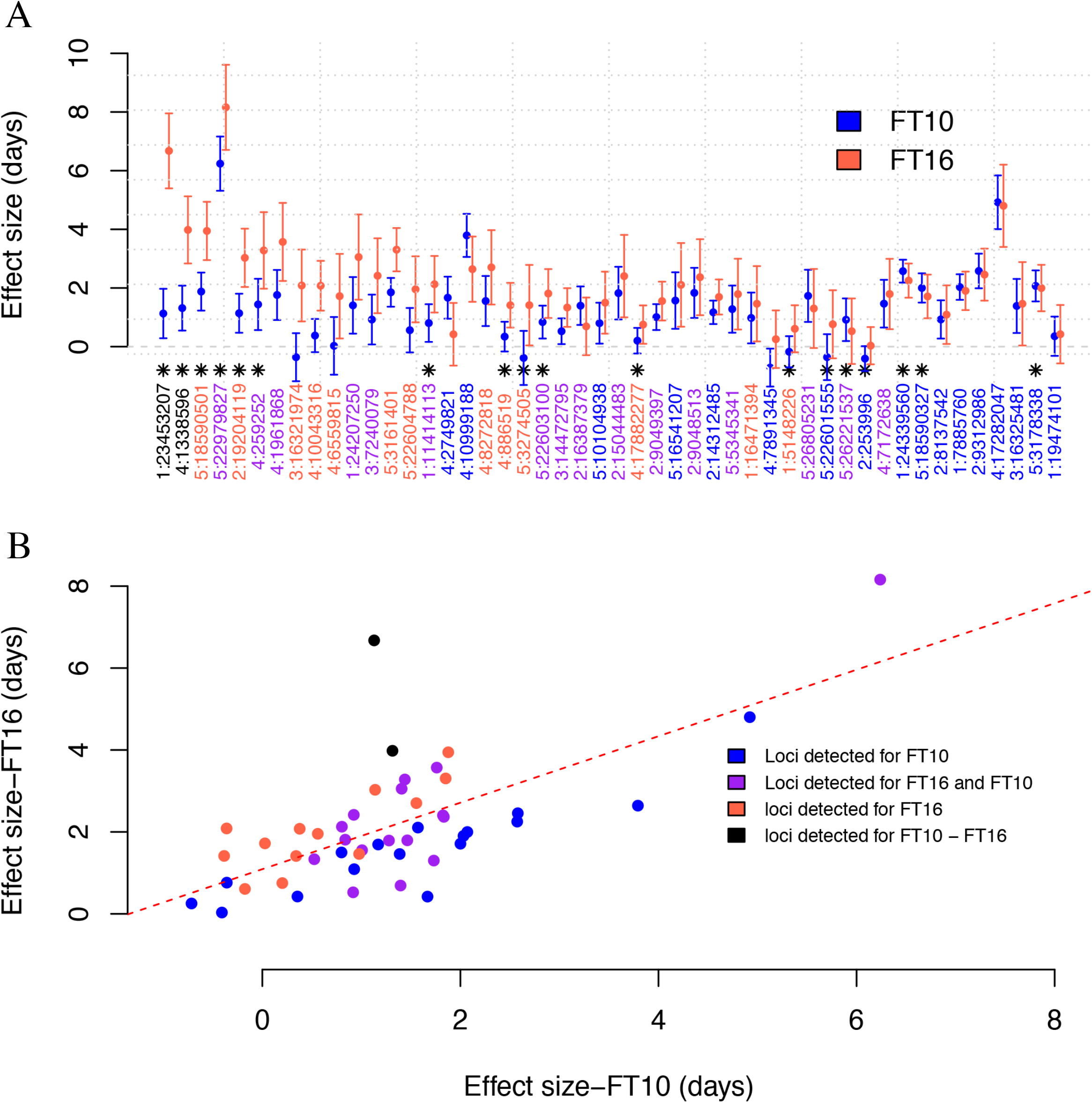
Additive effects of 48 loci on flowering time measured at 10 and 16 °C. **(A)** Additive genetic effects of 48 flowering time loci on FT10 (blue) and FT16 (tomato). The y-axis indicates the additive effects on flowering time and the x-axis the respective loci (SNPs) sorted by the differences in additive effects (FT16-FT10) from left to right. The estimates were obtained by fitting all 48 loci jointly in a mixed model. The bars represent standard errors of the effect estimates. The 18 loci with significant genotype by temperature interactions are indicated with a black star. **(B)** Correlation between the additive genetic effects for the 48 loci estimated for accessions grown at 10 and 16 °C. The colour of the dots indicates whether the locus is significant i) only for FT10 (blue), only for FT16 (tomato), for both FT10 and FT16 (purple), or the difference between the two (FT10-FT16; black).

### Distribution and effects of flowering time alleles in the worldwide *A. thaliana* population

To study the putative role of the 48 mapped flowering time loci in global and local adaptation, we evaluated how the alleles at these were distributed across the subpopulations identified in this dataset by *Alonso et al* [22]. The assumption in this analysis is that allele frequency differences between subpopulations across these loci, in its most extreme form the complete presence/absence of alleles, at least in part reflects past and ongoing selection leading to global adaptation along latitudinal or geographical clines, and local adaptations in individual subpopulations [33–35]. All 48 mapped loci were identified by having effects on flowering time at 10°C, 16°C, their difference or all, and we therefore consider all to be candidate adaptive loci as the temperature will vary both within and across the sampling sites of the accessions.

Consistent with the observation that the 48 mapped loci explained the majority of the additive genetic variance in flowering time, the overall pattern of early to late flowering times between the subpopulations in this dataset is captured well by the combined additive effects of these (Fig 3A/S4 Fig). Further, late flowering populations have accumulated both major and minor effect late flowering alleles across many loci (Fig 3B; late flowering alleles in red; effect sizes sorted from top to bottom). The late flowering north Swedish accessions are most extreme, with the highest frequencies at many loci. A visual inspection of the allele frequency patterns of the 48 loci across the populations (Fig 3B) suggests a greater overall similarity between populations from nearby geographic locations. This is analytically supported by a clustering analysis across the subpopulations based on the allele frequencies across these 48 loci (Fig 3C), and the pairwise correlations of the allele-frequencies between the populations (S5 Fig).

**Fig 3.**
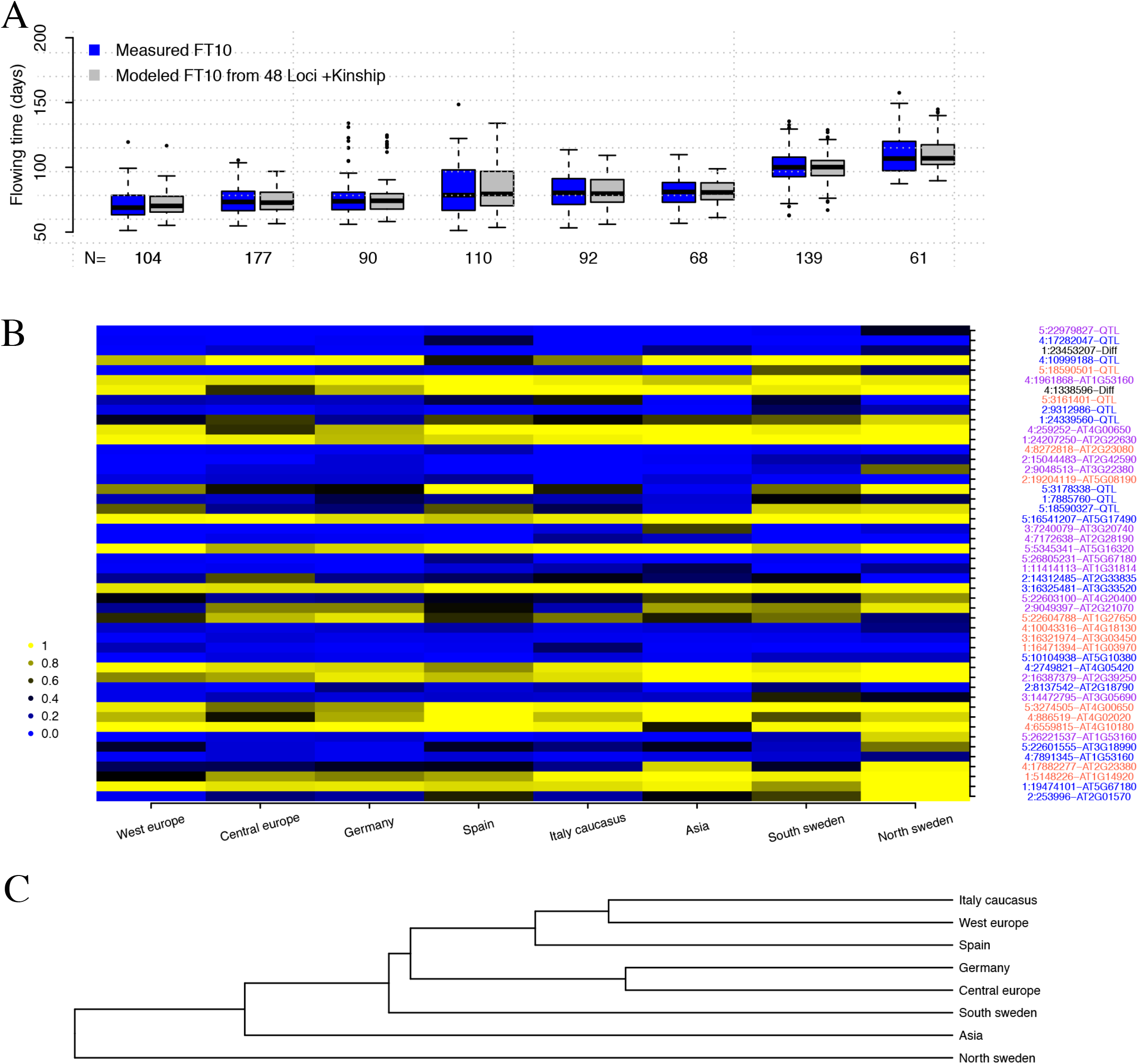
Flowering times, allele frequencies of flowering time associated loci and similarities of these between eight subpopulations of the *A. thaliana* 1,001-genomes collection. **(A)** Flowering times for the 8 subpopulations in this data defined by [22] are modelled well by the additive effects of the 48 flowering time loci and the kinship. **(B)** Distribution of allele frequencies for the 48 loci in the 8 subpopulations. Intensity of blue/yellow illustrates frequency of early/late-flowering alleles, respectively. The subpopulations are sorted from left to right by increasing average flowering time (columns) and the loci from top to bottom by decreasing additive effects on flowering time (rows). X:YYYYY-ZZZ represent a locus as Chromosome:Position-Type, where Position is in bp and Type is either “QTL”, “Diff” or “locus-id” indicating its detection by direct association to FT10, FT16 or (FT10-FT16) in a GWAS (“QTL”/”Diff”) or via a genetically regulated expression of the locus “locus-id” in the eQTL analysis. **(C)** Dendrogram showing how the 8 subpopulations cluster based on the allele frequency spectrum across the 48 associated loci.

For example, the accessions from Germany and Central Europe are closely related (Figs 3B and 3C), with an exception being the relatively late flowering southern Swedish accessions that overall were more similar to the earlier flowering Central European/German accessions than the late flowering north Swedish accessions.

For some of the 48 loci, both alleles are present (here defined as having MAF > 0.05) in all subpopulations, for example the loci on chromosomes 2 (16,387,379bp) and 4 (1,961,868bp) (Fig 2B; S6A/S6B Fig). For others, one of the alleles is either rare (MAF < 0.05) or absent in one or more of the subpopulations, for example chromosomes 2 (9,312,968bp), 4 (1,728,204bp) and 5 (22,979,827bp) (Fig 2B; S6C/S6E Fig). Both global and local allelic variations are thus likely to contribute to the natural flowering time variation across the worldwide range of *A. thaliana.* To explore this further, the 48 mapped flowering time loci were divided into two groups containing either those that are present across all populations (MAF > 0.05 in all populations; n = 23), or those that are rare or absent in at least one subpopulation (MAF < 0.05 in at least one population; n = 25). The choice of 0.05 as MAF cut-off is arbitrary but nevertheless serves the purpose of representing the spatial distribution of global vs primarily local alleles (Fig 3; Fig 5), and the results are overall similar also for MAF < 0.03 and MAF < 0.10 (results not shown).

Next, the contributions by global and local alleles to flowering time variation in the 8 subpopulations was evaluated. A mixed model, including the IBS kinship and the genotype of all 48 loci, was first fitted to FT10 and FT16 separately. Then, the estimated additive effects of these 48 alleles were used to calculate the contributions by local and global alleles, as defined above, to flowering time variation.

#### Contributions by global alleles to flowering time variation

For FT10, the IBS kinship captures the overall early to late flowering time pattern for subpopulations well and explicitly modelling the global alleles improves it further (Fig 4A). This illustrates that the polygenetic effect, together with the gradual allele frequency shifts across the loci with global alleles, are important for the overall pattern of flowering time differentiation between the subpopulations. The majority of the global alleles have moderate to small effects on flowering time (Fig 4A/B), relative to the local alleles (discussed further in section below), but together they contribute 21% of the phenotypic variance of the trait measured at both 10°C and 16°C. This analysis also shows that the global alleles have a larger contribution at 16°C than that to 10°C, again suggesting a role of genotype-by-temperature interaction for this trait (Fig 4A/B). The kinship is estimated to make a negative contribution to flowering times measured at 16°C in some subpopulations (Fig 4B), while the joint contribution by the global alleles is larger than at 10°C. One likely interpretation of this is that some of the global alleles have large overall effects at this temperature. They seems to also differ in effects across subpopulations and this genotype-by-environment interaction is captured by the kinship.

**Fig 4.**
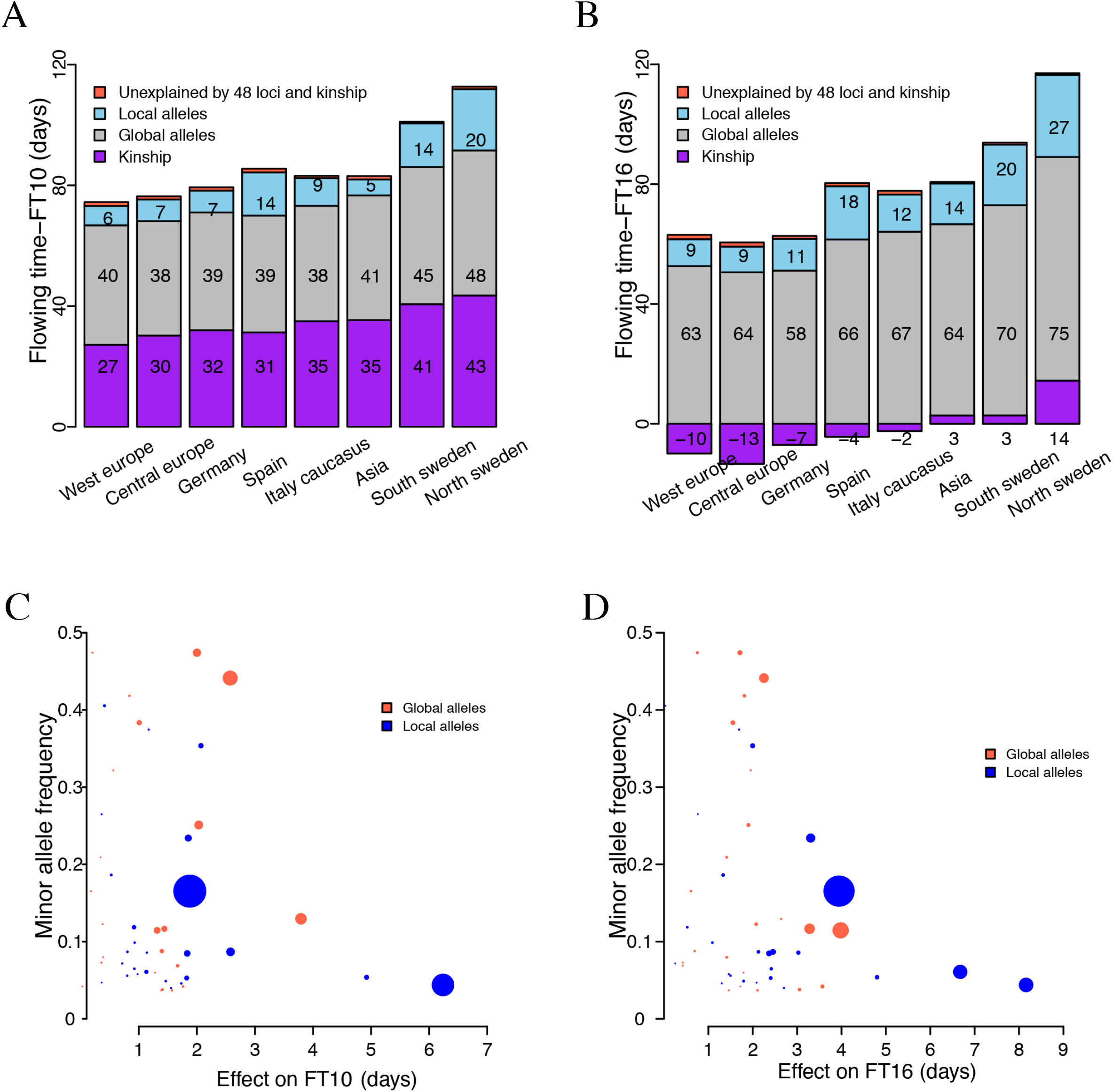
Modelling of the flowering times of the eight subpopulations in the 1,001-genomes *A. thaliana* collection and how the individual associated loci contribute. Experimentally measured and modelled flowering times in the 8 genetically defined subpopulations of the *A. thaliana* 1,001-genomes collection of accessions [22] grown at 10°C (**A**) and 16 °C (**B**). The height of the bars represents the experimentally observed mean flowering times for the accessions belonging to the sub population. The kinship (purple) and additive effects of globally segregating alleles (grey) capture the overall pattern of early to late flowering times between the sub populations well. The additive effects of the local alleles (blue) improve the fit to more extreme adaptations of some sub populations, in particular in Spain and Sweden. The differences between measured and modelled flowering times (tomato) are small at both temperatures. The modelled contributions by the global alleles are larger at 16°C than at 10°C due to their genotype-by-temperature interactions (Fig. 2A). The relationship between the variance contributed by each individual locus and its allele frequency/additive effect is illustrated for FT10 (**C**) and FT16 (**D**). The y-axis gives the minor allele frequency and the x-axis the estimated additive effect on flowering time in days. The size of each dot is proportional to the phenotypic variance explained by the locus. The colour of the dot indicates whether the minor allele is defined as a global (tomato) or local (blue) allele in the population.

#### Contribution by local alleles to flowering time variation

We next explored the loci with alleles that are rare or absent (MAF < 0.05) in at least one of the populations in more detail (Fig 5). Across these loci, the most apparent enrichment is for late flowering alleles in the Swedish populations (Fig 3B), where one or both of these populations contain near private late flowering alleles at high frequencies. These include the strongest late-flowering allele of all (chromosome 5 at 23 Mb; Fig 5), suggesting this large effect allele having emerged locally and been under strong selection here. A few other loci with relatively large effects on flowering were also predominantly present in one or a few subpopulations, but none were as striking as the aforementioned locus (Figs 3B and 5; S4 Fig).

**Fig 5.**
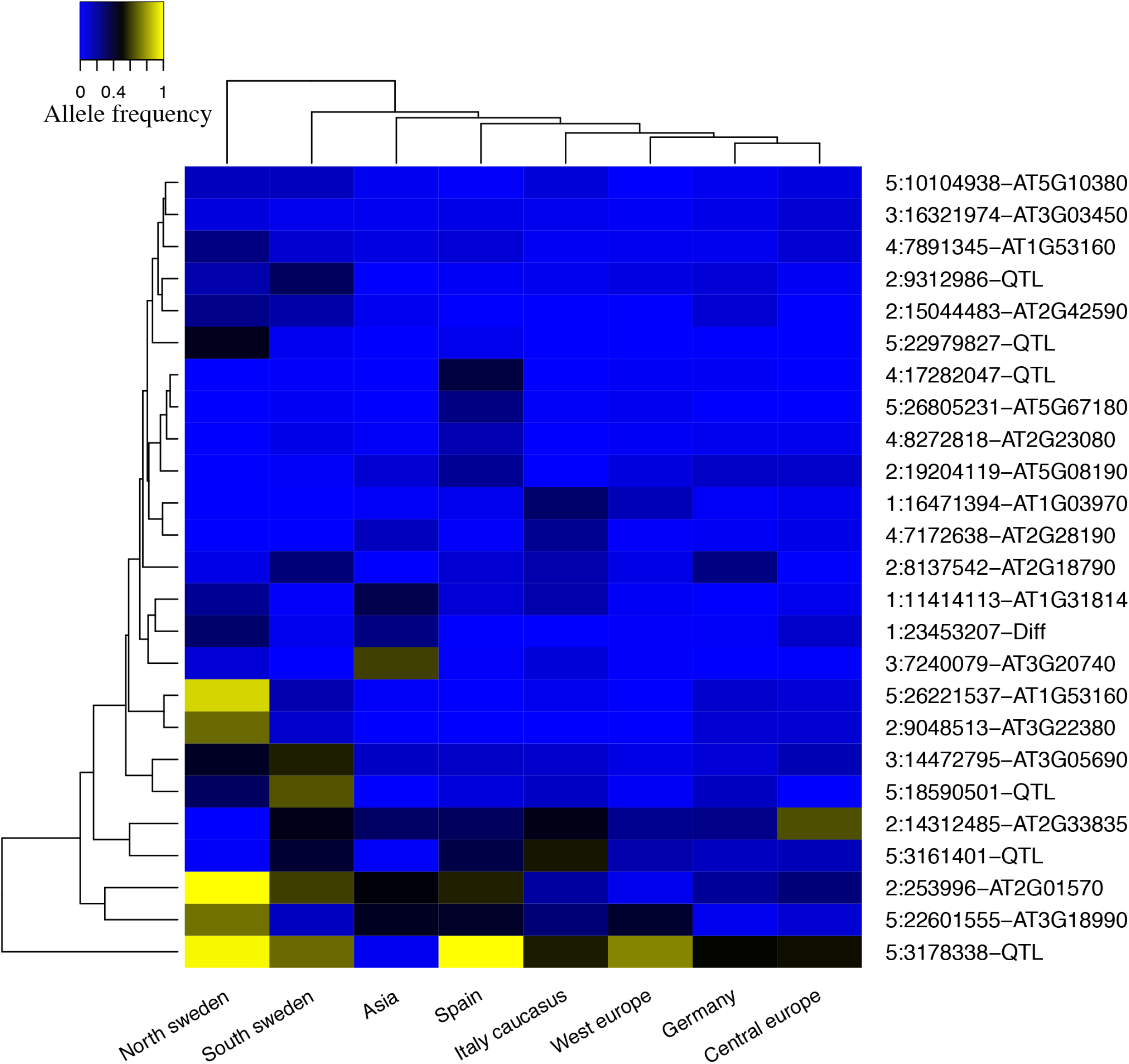
Allele frequencies for loci where one variant is rare (MAF < 0.05) in at least one of the eight 1,001-genomes *A. thaliana* subpopulations. Blue/yellow indicates high allele frequency for the early/late flowering alleles, respectively. The populations are clustered based on the overall similarity of the allele frequency pattern (left to right) and the loci based on the similarity of the allele frequencies in the subpopulations (top to bottom). X:YYYYY-ZZZ represent a locus as chromosome: position-type, where Position is in bp and Type is either “QTL”, “Diff” or “locus-id” indicating its detection by direct association to FT10, FT16 or (FT10-FT16) in a GWAS (“QTL”/”Diff”) or via a genetically regulated expression of the locus “locus-id” in the eQTL analysis.

The local alleles appears to primarily be contributing more extreme phenotypes in individual subpopulations (Fig 4A). In contrast to the global alleles, there is not only generally larger allele frequency differences between the populations for the local alleles, this group is also enriched for alleles with large effects that individually explain larger proportions of the phenotypic variance (Fig 4B). Together the local alleles contribute 45/42% of the phenotypic variance for FT10/FT16 in this population. The flowering time differences between the subpopulations thus result from shifts in the allele frequencies of global alleles (Fig 3B), with more extreme (for example northern Sweden) or variable (for example Spain) local phenotypes being due to private, or near private, local alleles (Fig 5).

### Long-range LD between flowering time loci

Long-range LD between loci is a possible sign of co-selection of them. To screen for such signatures, we calculated the pairwise LD (D’) between the 48 associated loci (Fig 6). In total, 67 pairs were in significant nominal long-range LD (P-value < 0.05; overall FDR for this set = 12%), where a few loci were in long-range LD with several other loci on either the same or different chromosomes (Fig 6). When a locus was in long-range LD with more than 4 other loci, the loci were defined as belonging to an LD-cluster.

**Fig 6.**
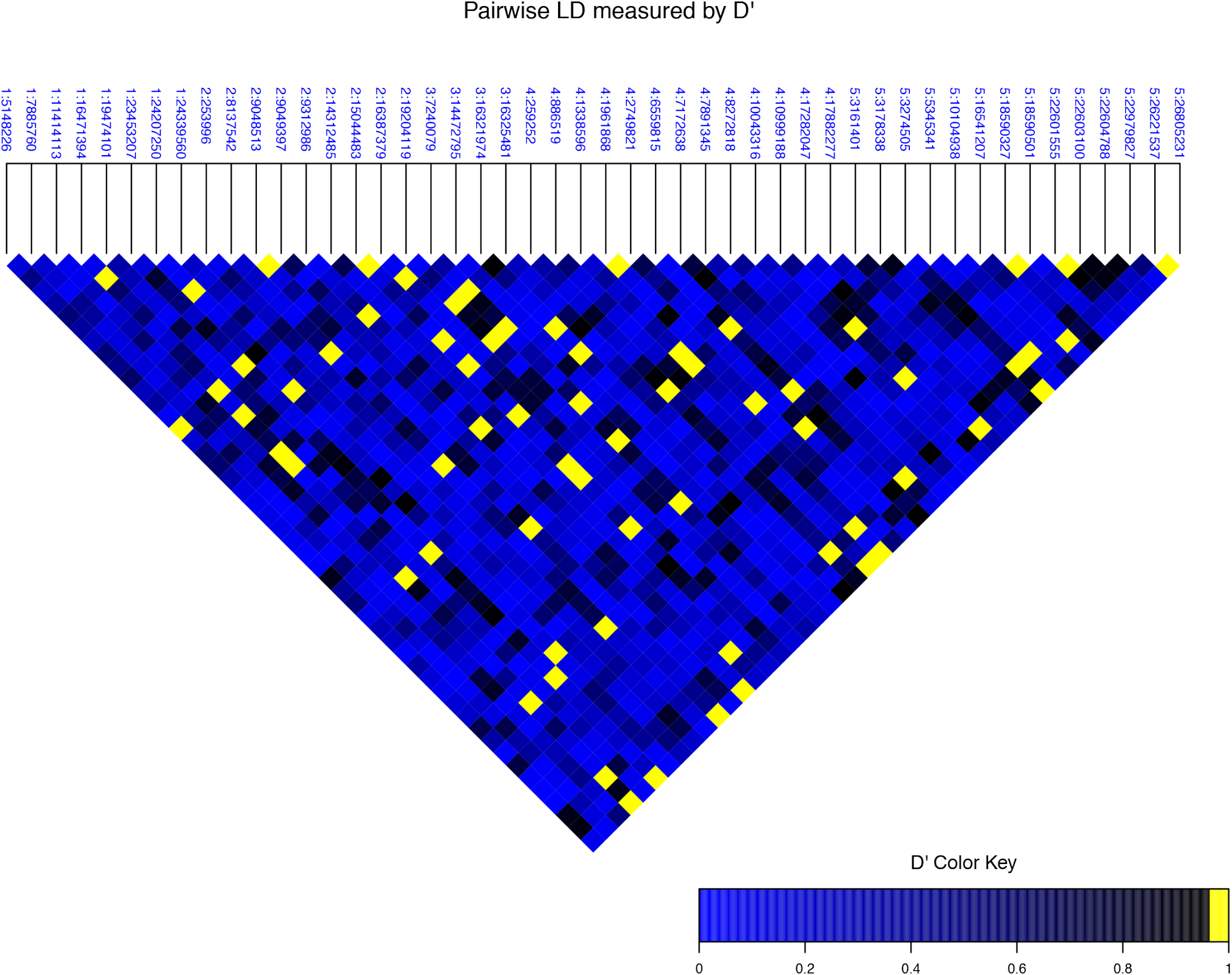
Pairwise linkage disequilibrium (LD) between the 48 flowering time-associated loci across the genome (D’). Values above the 0.99 quantile of a simulated null distribution are highlighted as yellow. The loci are labelled as chromosome: position on chromosome in bp.

We explored the five identified clusters in more detail. A literature review suggested functional connections between candidate genes in three of the five clusters. One cluster contained several genes in a miRNA mediated flowering time regulation pathway, suggesting a functional connection to the FLC-CO/FT module [36–41] (Table 1). Two clusters contained genes whose function are connected to the *FRI-FLC* module, one of these possibly through vernalization [42–44]. The minor alleles of the five loci that via their long-range LD defined these clusters were enriched in specific geographic regions. The one allele on chromosome 2 (15,044,483bp) was found primarily in Sweden, the one on chromosome 1 (16,471,394bp) in Italy Caucasus, the one on chromosome 3 (7,240,079bp) in Asia, the one on chromosome 4 (17,282,047bp) in Spain and the one on chromosome 5 (22,979,827bp) in north Sweden (Fig 4B). Together this suggests possible contributions from these pathways to local adaptation in these geographic regions.

**Table 1.**
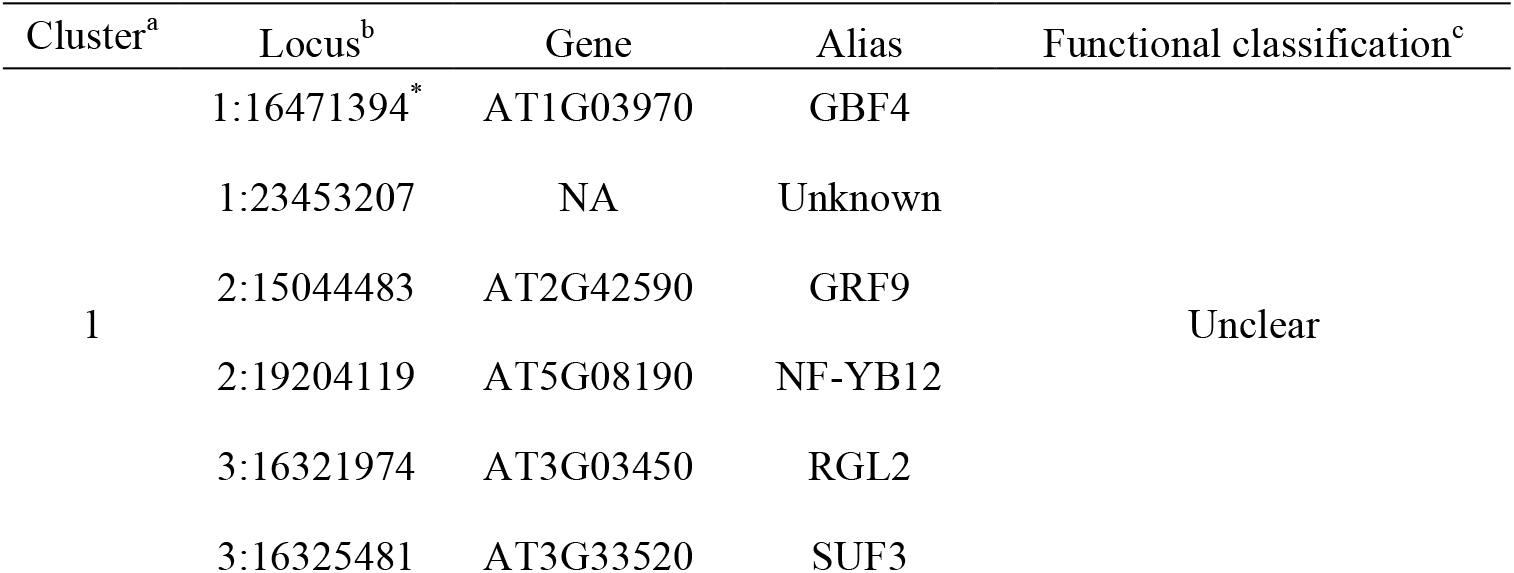
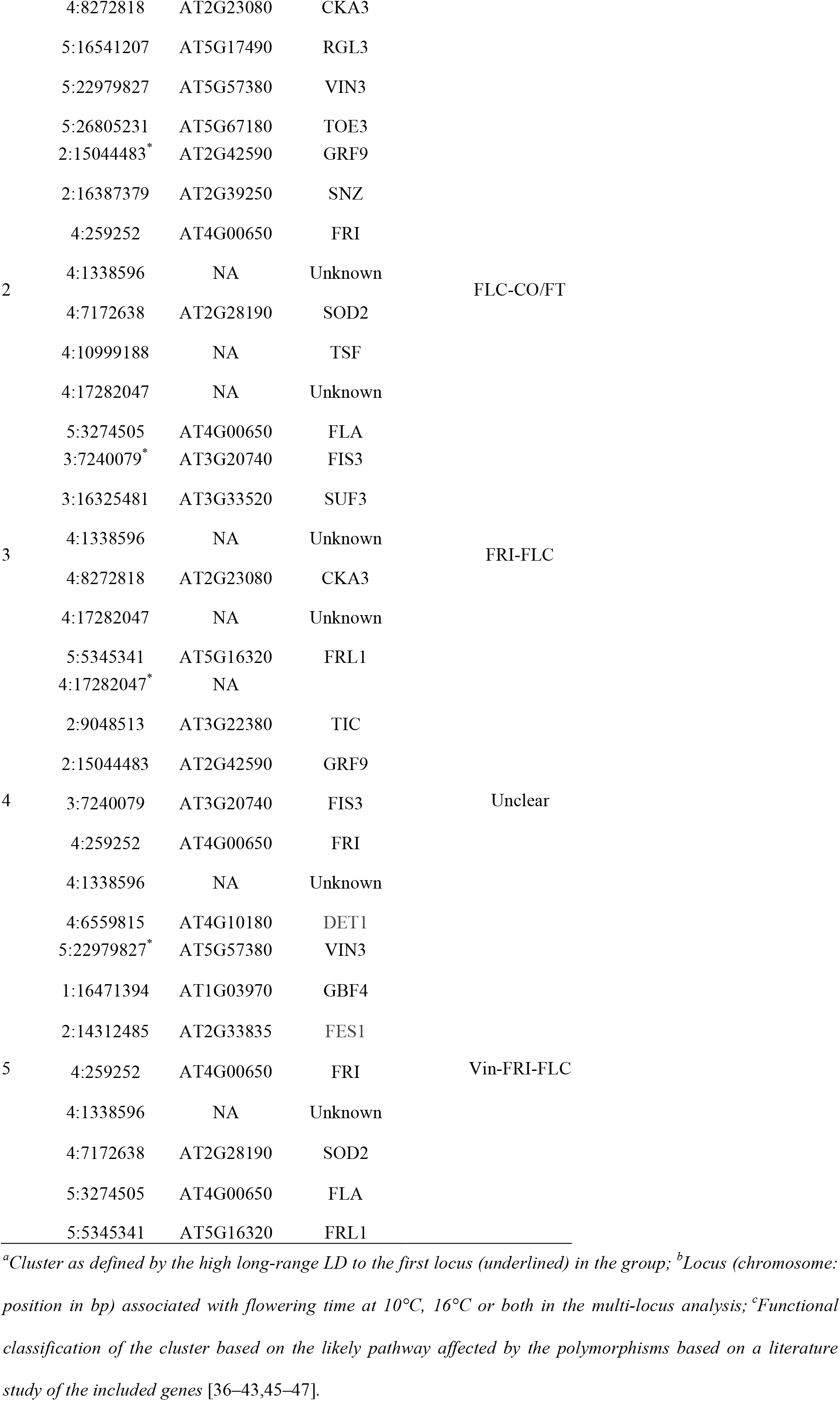
Long-range LD defined clusters of genes with assigned functional classification

## Discussion

In an earlier study, we mapped the polygenetic basis of flowering time variation in the latest public release of data from the 1,001 genomes *A. thaliana* project [22,26]. In this analysis, 33/29 loci were reported to be associated with flowering time variation measured at 10/16°C and they together explained 66/58% of the additive genetic variation[26]. These loci were mapped using a two-step polygenic analysis method. First, loci with individual genome-wide significant associations either directly to measured flowering times, or to the expression of experimentally known flowering time genes, were identified as candidate genes for the polygenic analysis in this population. Then, a joint analysis across all these candidate loci was performed for each trait separately using a backward elimination analysis, to infer a polygenic architecture of the trait at a 15% false discovery rate significance threshold. At this significance threshold, a few of the 33/29 loci associated with FT10/FT16 are expected to be false positives. Here, we continued this work by treating flowering time as one trait measured independently for the same genotypes at 10°C and 16 °C. The focus was to evaluate the joint contributions by all mapped flowering time loci to the genetics of this trait and their possible role in global and local adaptation to the different ecological conditions across the worldwide range of this plant. We found that the groups of loci that were individually significant in one growth temperature, but not the other, made significant joint contributions to flowering time in both conditions. Fitting all 48 loci together in a multi-locus linear model, they explain most (79%/75% for FT10/FT16) of the additive genetic variance in flowering times at these growth temperatures. As most of the loci are likely to be true associations, we performed an in-depth evaluation of how this set of loci contribute to the global flowering time variation in *A. thaliana* to provide valuable complementary information to earlier studies that have focused primarily on loci with strong individual associations[23,25,48,49].

It is generally difficult to study the genetic basis of highly polygenic traits in natural populations due to, for example, the small contributions made by individual loci to trait variation and the confounding of adaptive alleles with population structure. This is likely one of the reasons why standard genome-wide association mapping approaches have only been able to reveal a small number of flowering time loci in this or related populations [22–24]. By approaching flowering time as a polygenic complex trait, and making use of its measures in two temperatures, this study could shed lights on how allele frequency changes across many loci, as well as genotype by temperature interactions, contribute to the genetic basis of global and local trait variation.

### The role of global and local alleles in shaping worldwide flowering time variation

For many of the polymorphic flowering time loci, both alleles were present in all subpopulations of the worldwide population, albeit at different frequencies. Together these contributed to the overall pattern of early to late flowering in the geographically divided subpopulations. Polygenic global adaptation via small allele frequency shifts across many loci thus appears to be a possible contributor to the variation of this trait in *A. thaliana.*

For a number of the associated loci, one of the alleles was absent or present at low frequency (MAF < 0.05) in one or more of the subpopulations. Several of these local alleles had strong effects on flowering time, and these loci made important contributions to the more extreme flowering time phenotypes of individual subpopulations. Alleles with large effects on adaptive traits have earlier been reported to be present in local populations of *Arabidopsis thaliana* [5,7,23,50]. Our finding that many such alleles are present in locally in the worldwide population, is coherent with earlier speculations that locally adaptive alleles for flowering time variation in *Arabidopsis thaliana* are likely to have large effects [51]. Those local populations appear to reach their respective flowering times via the effects of alleles that are primarily present in some populations suggests that different combinations of alleles are likely explaining adaptation to different local environments.

We evaluated the distribution of many flowering time alleles across the native range of the species. The results are interpreted under the assumption that the alleles associated with flowering time are very likely to have an adaptive value and that the observed patterns of allele frequency differences between populations, to large extent, represent local and global adaptations. Other explanations are, however, also possible as similar patterns could emerge due to, for example, an interplay of drift, historical contingency and selection. A few observations are particularly valuable when interpreting the results. First, the consistency in which loci are mapped, and their patterns of effects when the plants are grown in two different growth temperatures, suggests that the genetic architecture of flowering time is overall stable. Hence, even though the individual effects of many loci are small, making it difficult to assess the selective pressure on each locus in nature, polygenic adaptation acting over time on the many alleles that are present globally in *A.thaliana* appears like a possible route to adaptation. The finding that stronger local alleles complements the global polygenic adaptation, likely through a more rapid increase in allele-frequencies due to stronger selection for locally favourable phenotypes, is also consistent with earlier reports in which a few flowering time QTL were found to be correlation with latitude [48, 52], indicating a potential role of spatial adaptation. That the same process might have been important also for other traits is suggested by the results of another recent study on drought adaptation reporting a polygenic genetic architecture and accompanying geographical enrichment of QTL[53]. Another possible explanation for the finding of, more or less, local alleles in several subpopulations is that the plants are naturally restricted to its native range over their lifespan and that self-pollination in *Arabidopsis thaliana* slows down the long distance spreading of new genetic variants. Hence, newly emerged alleles with large beneficial adaptive effects can quickly sweep to high frequency in local populations, but long distance spreading of those alleles will take longer in this selfing plant, although it is known to happen due to for example human or animal activities [54]. A possible example of such long distance spreading of genetic variants observed in this data, is the large effect allele identified on chromosome 5, 23 Mb. Here, the strong allele is present at high frequency in northern Sweden and in a relict population in Spain (S4F Fig). These populations are geographically distant, and it appears a likely result from lineage mixing between relict and northern populations as indicated in an earlier study of this dataset [22].

Two late-flowering alleles, with intermediate to high effect, are private (or nearly so) to the Spanish population (Fig 4). We note that this population has a significantly higher within-population variation in flowering time (P = 1.1 × 10^−6^; Brown-Forsythe test) than the other populations (Fig 2A). The reason for this is not known, but a possible explanation could be that these strong alleles have contributed to specific local adaptations in this subpopulation. Further studies of these effects would be valuable to evaluate the possible contributions of these alleles to local adaptations.

### Possible contribution by co-evolution of functionally related loci in local populations

Four of the flowering time associated loci were in long-range linkage-disequilibrium (D’) with several other mapped loci. These alleles were enriched in relatively distinct geographic regions, suggesting that they have emerged and spread locally making a confounding with population structure a possible explanation for this observation. We, however, consider this unlikely as they were detected in a statistical multi-locus analysis accounting for population-structure and hence each of them captures flowering time variation that could not be explained by the other loci. Further, as in three of the five cases the loci involve candidate genes in related biological pathways. An intriguing alternative interpretation is that the LD instead result from co-selection of multiple functionally connected loci. Further work is, however, needed to confirm this hypothesis.

### Genotype-by-temperature interactions and their possible implications for adaptation

An earlier study reported genotype by environment interactions in the Swedish *A. thaliana* populations that were part of the dataset analysed in this study [30]. Here, we explored the possible role of genotype-by-temperature interactions for in total 48 flowering time loci associated with this trait in the entire 1,001-genomes collection of worldwide accessions grown at both 10°C and 16°C. Our conceptual approach to study these interactions was slightly different to the one used before. First, flowering time was considered as a single complex trait and all loci associated with this trait, regardless of growth temperature, was considered as possible loci involved in genotype-by-temperature interactions. Next, the effects of all the 48 associated loci on the flowering times measured at 10/16°C were estimated. The allelic effects in the two growth temperatures were compared, and possible genotype-by-temperature interactions inferred based on observed differences in effects between the two temperatures. Given the high phenotypic correlation between flowering times of accessions grown in these different temperatures (Pearson correlation=0.88), we expected the effects to be overall similar between environments. However, for ∼1/3 of the mapped loci (18 of 48 loci; Fig 6), the effects on flowering time changed with temperature. These results suggest that the genetic effects of many loci in the genetic architecture of this polygenic trait are dependent on one or more environmental factors, as exemplified by temperature here. As studies aiming to detect complex trait loci, for example using a GWAS approach, are generally limited in their ability to assess the effects of loci across multiple environments a significant fraction of loci important in the variable conditions under which plants live in nature might be missed. More studies, for example to identify seasonally sensitive QTL in the field using GWA analysis[48], will therefore be of great value for our understanding of the genetic of such traits and their potential role in adaptation.

The potential adaptive values of the genotype-by-temperature interactions identified here is difficult to assess. We observed a geographical enrichment of loci involved in such interactions, that have larger effects at 16°C, in the northern Swedish populations, where flowering time is increased at higher temperature (S7 Fig). Further investigations uncovered significant pairwise correlations between the joint effects of these loci on flower time differences between temperatures (FT10-FT16) and within year temperature fluctuation (temperature difference between hottest and coldest day; S8 Fig). It is therefore possible that these loci are of adaptive value in environments where the temperature fluctuates drastically, and where having alleles that can shorten flowering time in response to cold temperatures would be beneficial. However, further studies are needed to evaluate this hypothesis thoroughly.

### Concluding remarks

Our study demonstrates that the variation in flowering times between earlier defined geographically and genetically separated *A. thaliana* subpopulations is likely due to allelic variations in many loci. At some of these loci both alleles are present across the entire range of the species, whereas others are only present in some of the subpopulations. Across these loci, the allele frequency shifts were often more gradual at loci with global present alleles. Larger differences in allele frequencies were observed for several loci with alleles of relatively large effects, and higher allele frequencies were then found in local populations with more extreme phenotypes. This suggests that flowering time adaptation results via a combination of global polygenic adaptation across many loci, complemented by local adaptation via the effects of stronger alleles with a more regional distribution. The results also indicate that local phenotypes might also result from co-evolution of multiple polymorphisms in biological pathways. We identified genotype-by-temperature interactions involving 18 flowering time loci, and propose a hypothesis that these might be adaptive in environments with large temperature fluctuations. However, further investigations are needed to validate this hypothesis. Overall, this study provides new insights to the genetic basis of flowering time in *A. thaliana* and propose possible explanations for how this self-pollinating plant has been able to colonize and adapt to such a wide range of ecological habitats around the world.

## Materials and Methods

### Data

All phenotype, genome-sequence and subpopulation information are publicly available as part of the *Arabidopsis thaliana* 1,001-genomes project [22, 55]. Flowering times measured at 10°C and 16°C in the greenhouse were downloaded from [56, 57]. The subpopulation classifications of the accessions were downloaded from http://1001genomes.org/tables/1001genomes-accessions.html. The imputed whole genome SNP data matrix was downloaded from http://1001genomes.org/data/GMIMPI/releases/v3.1/SNP_matrix_imputed_hdf5/1001_SNP_MATRIX.tar.gz. We filtered the SNP data for minor allele frequency and only retained loci with MAF > 0.03. SNP markers were pruned to remove loci in pairwise LD of r^2^ > 0.99. In total, 1,396,438 SNPs on 1,004 individuals remained. The 33/29 loci associated with flowering times at 10/16°C were identified in a previous study[26]. These loci were inferred using a two-step polygenic mapping method. First, candidate flowering-time loci were identified as those with either i) individual genome-wide significant associations to flowering time or ii) that altered the expression of genes that are experimentally known to affect flowering. Then, these candidate loci were analysed jointly for associations to flowering-times measured at 10°C and 16°C using a backward elimination based multi-locus mapping approach to infer the polygenic architecture of flowering time at a 15% false discovery rate significance level [26].

### Detecting loci involved in genotype by environment interactions

Two methods were used to test for QTL by environment interaction. The first was a genome wide scan using the mean flowering time difference between 10°C and 16°C as phenotype. In the analysis, a forward-selection based approach was employed where a linear mixed model *ȳ*_FT10_ – *ȳ*_FT13_ = Xβ + Zu + e (1) was iteratively fitted to the SNPs across the genome with significant SNPs from earlier rounds added as covariates in the model. This model has earlier been shown to have equivalent power with a bi-variate analysis for detecting loci with increased/decreased effects across environments[58]. Here, *ȳ*_FT10_ – *ȳ*_FT16_ is the mean flowering time difference between 10°C and 16°C for an individual accession. *X* is the design matrix including a column vector of 1’s for the population mean and one or more additional column vectors with the indicator regression variables for the homozygous genotypes at the SNP or SNPs included in the model (coded as 0/2, respectively). *Z* is matrix obtained by singular value decomposition of the *G matrix* (*ZZ*^T^ = *G*), where *G* is the genomic kinship matrix estimated from the whole genome marker set using the *ibs* function in the *GenABEL* R-package [59]. The significance-threshold used in the analysis was Bonferroni corrected for ∼1.4 million tested SNPs (-log10(p-value)=7.33).

Second, the loci associated with flowering-time in the earlier polygenic analysis were explored for genotype by environment interactions. Two models were fitted to the data:

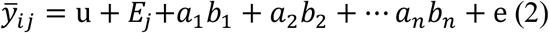

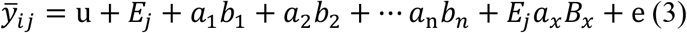

In both models, *ȳ_ij_* is the mean flowering time among replicates of individual *i* phenotyped in environment *j* where *j* here take the values 1 and 2 for FT10 and FT16, respectively; *a_x_* is the genotype at QTL x coded as 0 and 2 for the homozygous minor and major allele genotypes; *b* are the corresponding estimated effect sizes; *u* is the population mean and *E_j_* is the effect of environment on flowering time. Model 3 includes an interaction term between one of the QTL and the environment *E*, and was fitted for each of the QTL in turn. The significance of each QTL by environment interaction was assessed using a likelihood ratio test between model 2 and model 3. Population structure was accounted for by the simultaneous fitting of the 48 loci. The analyses were performed using customised R scripts [60].

### Evaluating how groups of loci contribute to flowering time variation

Several of the mapped loci are only associated with flowering-time in one of the growth temperatures. We tested whether the flowering-time loci that were not significantly associated individually in one of the temperatures made a significant joint contribution by comparing the fit of the following models to the data (models 4 and 5) using a likelihood ratio test.

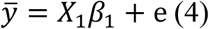

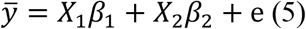

Here, *ȳ*, and e are defined as for model 1. Contributions from the individually significant/nonsignificant loci in the tested temperature were modelled as *X*_1_*β*_1_/*X*_2_*β*_2_, respectively. *X_1_* is a similar design matrix as *X* in model 1, which includes a column vector of 1’s for the population mean and one or more additional column vectors with the indicator regression variables for the homozygous genotypes at the SNP or SNPs included in the model (coded as 0/2, respectively). *X*_2_ is a second design matix only including the genotype for additional loci unique to FT10 or FT16. *β*_1_/*β*_2_ are the corresponding effect sizes. A likelihood ratio test was used to compare the fit of the two models using the *lrtest* function in R package *lmtest* [61].

### Clustering of subpopulations based on the allele frequencies of flowering time associated loci

Euclidean distances between the 8 subpopulations in the 1,001-genomes dataset defined in [22] were calculated using the *dist* function in R based on the allele frequencies at the 48 loci associated with the flowering time at 10°C or 16°C in this dataset. The subpopulations were then clustered using the hierarchical clustering function *hclust* in R.

### Estimating the joint, local and global effects of all associated flowering-time loci

The effects of all loci associated with flowering-time in 10°C, 16°C or both temperatures (as in model 4) were estimated simultaneously using the *hglm* function in *hglm* R-package [62].

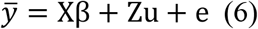

All parameters are the same as in model 1, except *X* in which a column vector of 1s is included to model the population mean and then *n* genotype columns (where the genotype for an individual at a locus is coded as 0 and 2 as above) for each of the *n* mapped flowering-time loci. The fitted values from model 6 were then used to model the contributions by global and local alleles to the flowering time. Contributions from the local alleles were estimated by *X*_1_*β*_1_, where *X*_1_ is the genotype matrix of the local alleles and *β*_1_ is the corresponding estimated effects from model (6). Contributions from the global alleles were estimated by *X*_2_*β*_2_, where *X*_2_ is the genotype matrix of the local alleles and *β*_2_ is the corresponding estimated effects from model (6). Contributions from the kinship were then estimated by subtracting *X_1_β_1_* and *X*_2_*β*_2_, from the modelled phenotype. Then, estimates for each subpopulation were obtained by averaging the individual estimates obtained for the accessions in it.

### Testing for significant long-range Linkage Disequilibrium

An empirical significance threshold was derived to test for significant long-range LD between the flowering-time loci associated with flowering time at 10°C, 16°C or both. An empirical null distribution was obtained using 1,000 simulations, where in each the same number loci as detected in the association analysis were simulated with allele frequencies being the same as those of the associated loci. In each simulated dataset, all pairwise LD (D’) values were calculated and saved to generate a null distribution for the significance test. We used the 0.99 quantile (corresponding to a D’ value of 0.96) as cut-off in our analyses. The FDR for the result was calculated as the ratio between the expected number and observed numbers of significant pairs (chose (*n*,2)*0.01), where *n* is the number of associated loci.

### Visualization of the results

Figs 3 and 5 were created using the *heatmap.2* function in *gplots* package in R [63]. Fig 6 was created using the *LDheatmap* function in the *LDheatmap* package in R [64]. Fig S3 was made using the R package *maptools* [65]. All other figures were created using custom R-scripts [60].

---

Supplementary information for **“Explorations of the polygenic genetic architecture of flowering time in the worldwide *Arabidopsis thaliana* population** “ by Zan and Carlborg.

**S1 Fig.**
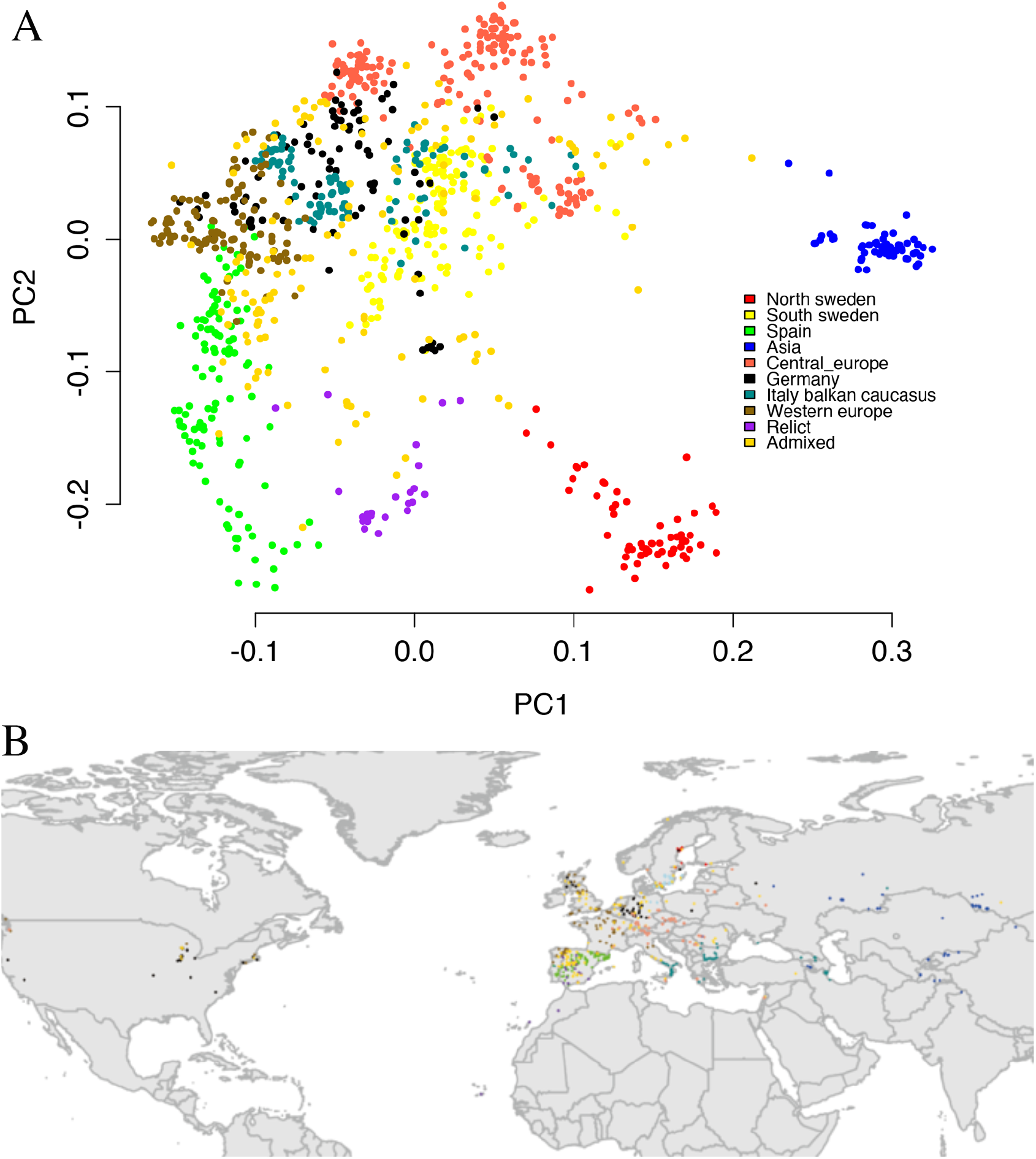
Population structure and sampling locations for the accessions in the 1,001-genomes *A. thaliana* collection. **(A)** Population structure indicated by the first two principal components of the IBS (Identity By State) kinship matrix with the subpopulations defined by [22] highlighted in colour. **(B)** Sampling locations of the analysed accessions.

**S2 Fig.**
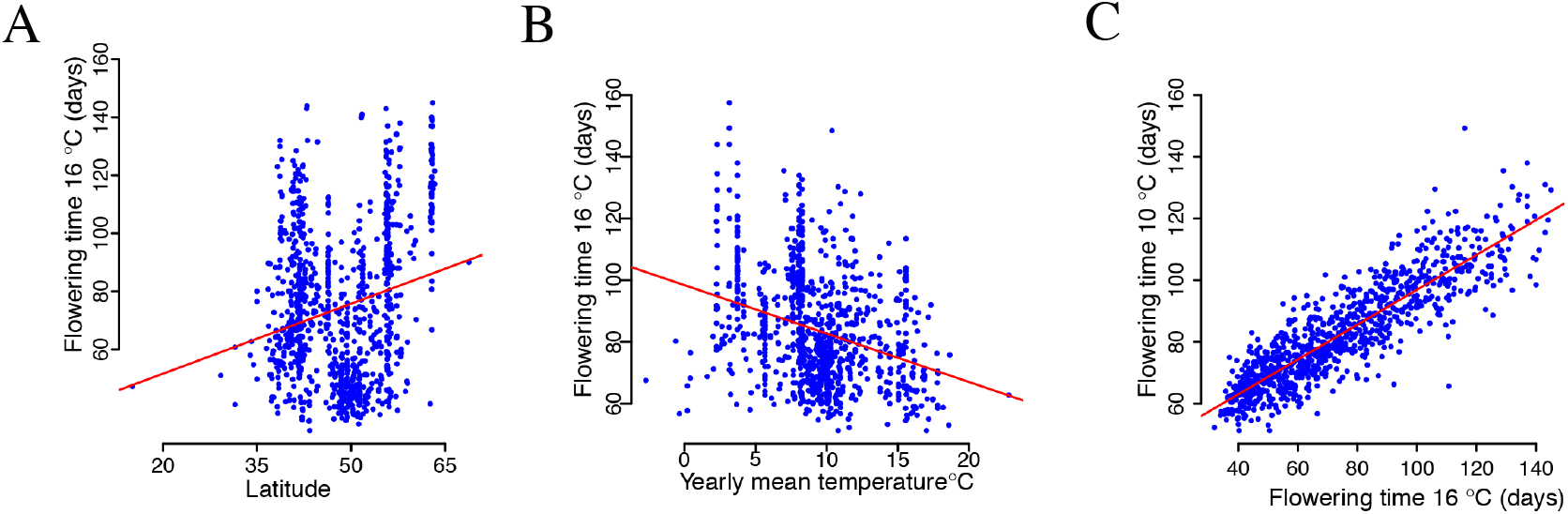
Geographical differentiation, and correlations, of the flowering times for the accessions of the 1,001-genomes *A. thaliana* collection grown at 10 and 16°C. Flowering times measured for accessions grown at 16°C in the green house are significantly correlated both with latitude (**A**) and temperature (**B**) at the sampling sites for the 1,004 analysed accessions from the 1,001-genomes *A. thaliana* collection [22]. (**C**) There is a high correlation between flowing times measured for accessions grown at 10 and 16°C.

**S3 Fig.**
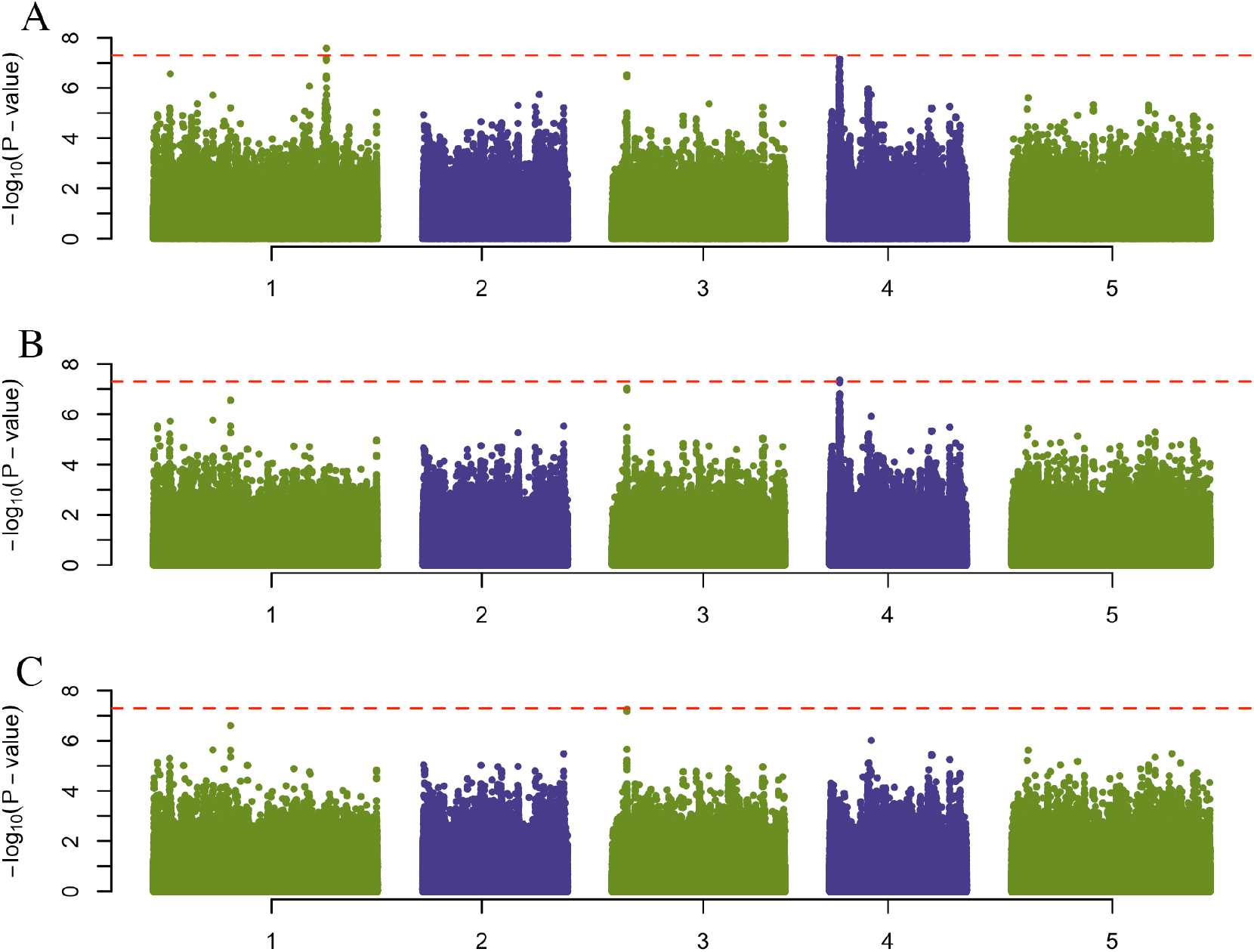
Results from the GWAS analysis using the differences in flowering times at 10 and 16°C for the accessions of the 1,001-genomes *A. thaliana* collection. (**A**) Manhattan plot for a GWAS using FT10-FT16 as phenotype where one locus on Chromosome 1 at 23,453,207 bp reached genome wide significance. (**B**) Manhattan plot for a GWAS using FT10-FT16 as phenotype with the locus identified in (**A**) as covariate, where a second locus on Chromosome 4 at 1,338,596 bp reached genome wide significance. (**C**) Manhattan plot for a GWAS using FT10-FT16 as phenotype with the 2 loci identified in (**A**) and (**B**) as covariates, where no additional locus was detected.

**S4 Fig.**
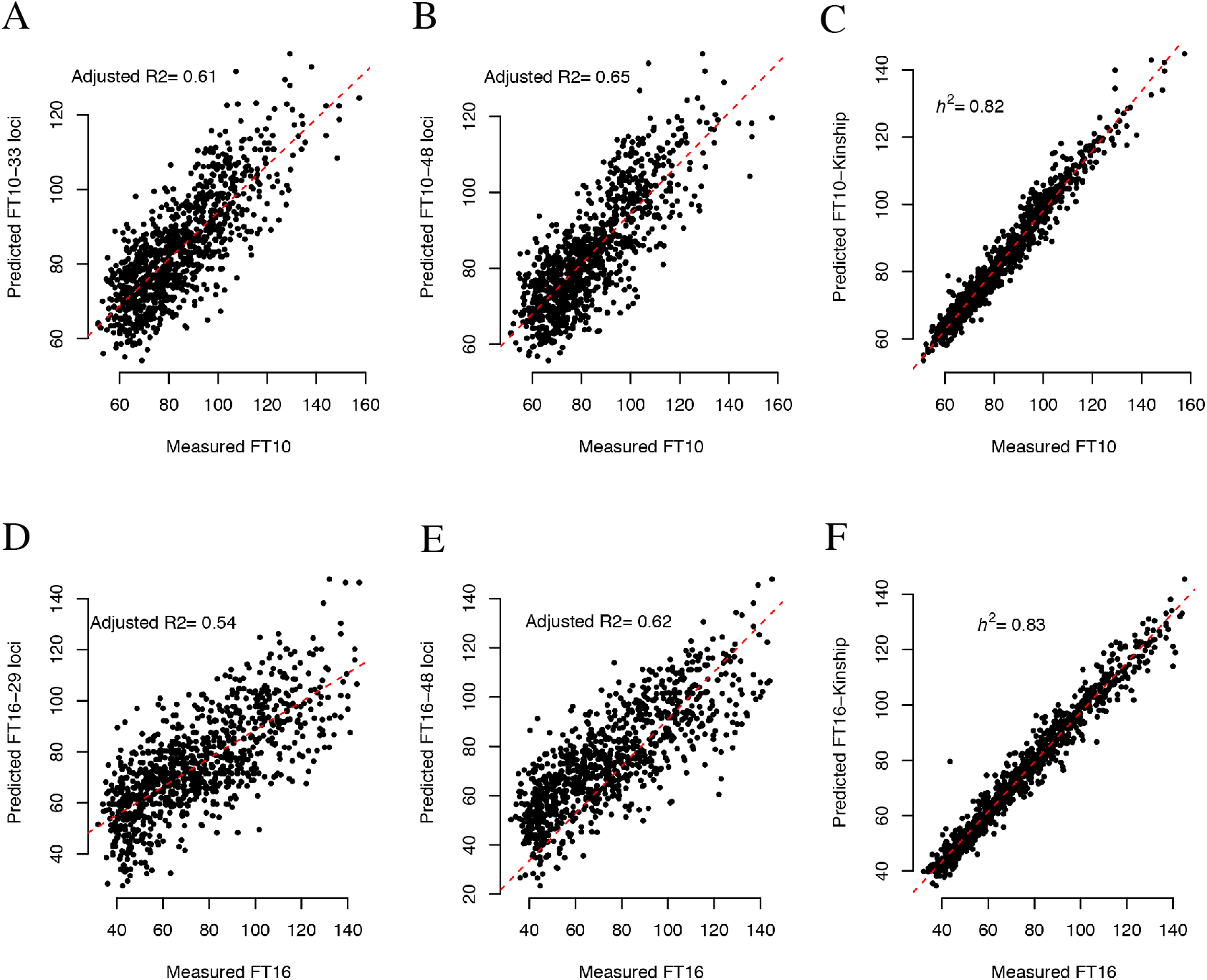
Modelled and experimentally measured flowering times for the accessions of the 1,001-genomes *A. thaliana* collection. Modelled phenotypes for the 1,001-genomes accessions using the 33 (29) loci detected in the independent analyses of FT10/FT16 and the experimentally measured phenotypes are illustrated in (**A**) and (**D**). Modelled flowering times using all 48 flowering time loci and measured phenotypes are illustrated in (**B**) and (**E**). Modelled flowering times from kinship and measured phenotypes for FT10/FT16 are illustrated in (**C**) and (**F**).

**S5 Fig.**
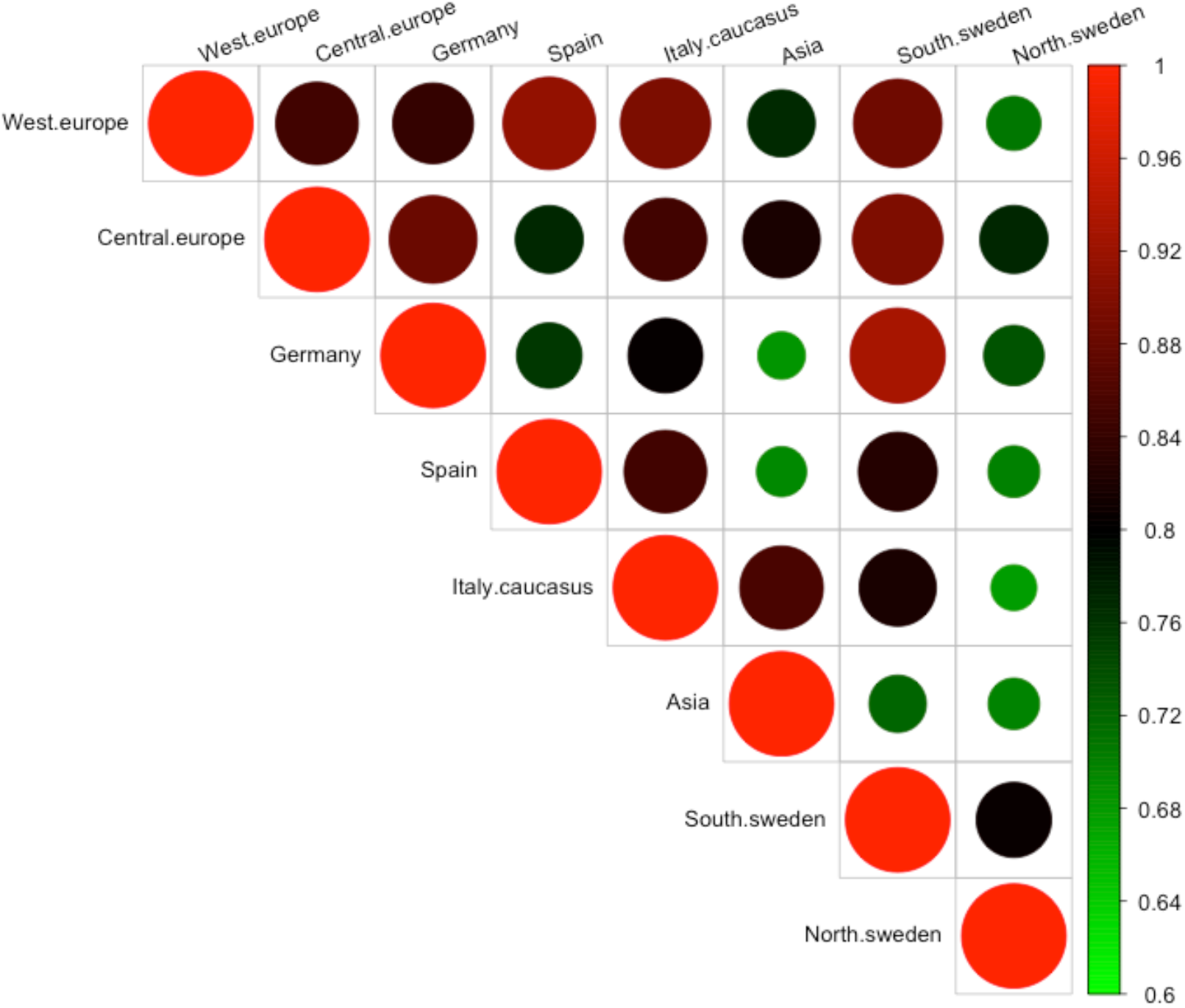
Illustrations of the overall similarities in the allele frequency spectrum across the sub populations as Spearman correlation between the allele-frequencies across the 48 associated loci

**S6 Fig.**
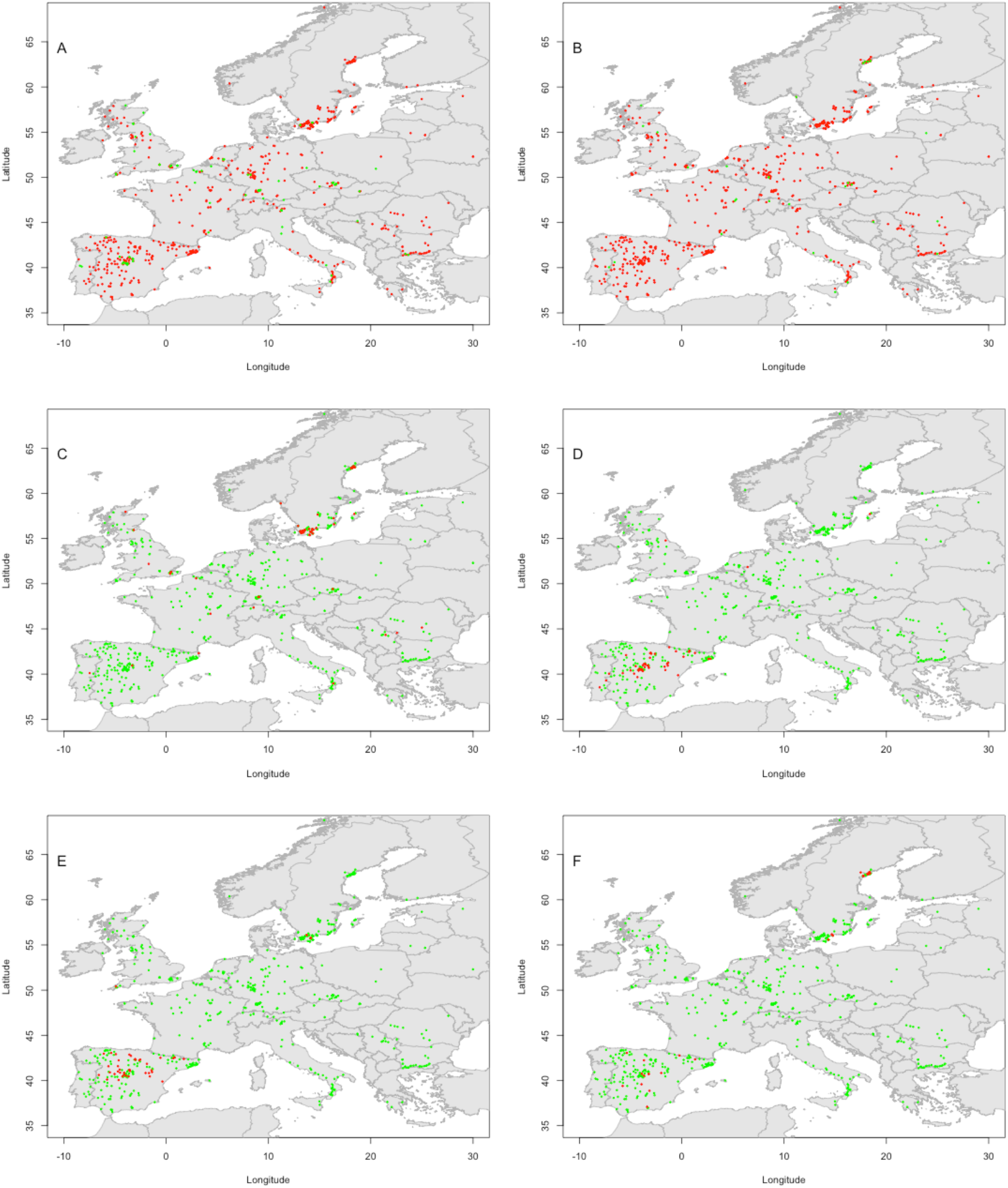
The geographical distribution of alleles at 6 selected loci associated with flowering-time at 10°C (FT10). The five panels illustrates the geographical distributions for the six associated loci at Chromosome 2:16,387,379 bp (**A**), Chromosome 4:1,961,868 bp (**B**), Chromosome 2: 9,312,968 bp (**C**), Chromosome 4:1,728,204 bp (**D**), Chromosome 5:26,805,231bp (**E**) and Chromosome 5:2,297,9827 bp (**F**). Red/green colours indicate the allele that increases/decreases FT10, respectively.

**S7 Fig.**
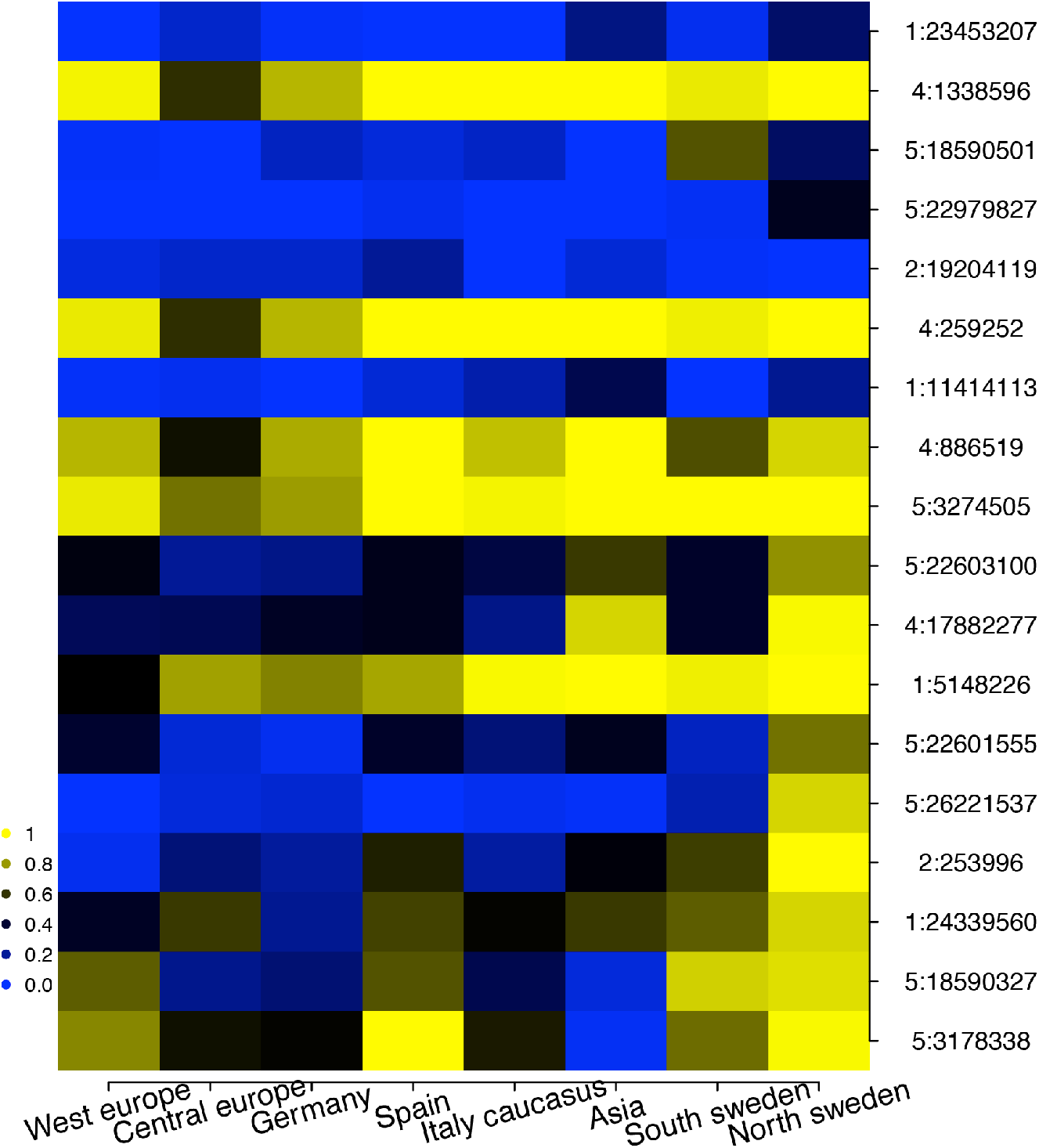
Allele frequencies for the 18 loci showing significant genotype by temperature interactions across the 8 sub populations of the 1,001-genomes collection. Blue/yellow indicates high allele frequency for early/late flowering alleles, respectively. The populations are sorted based on early to late flowering (left to right) and large to small additive effects on the flowering time difference (FT10-FT16; top to bottom). Genotype by temperature interacting alleles that have a larger effects on FT16 than on FT10 are enriched in the North Sweden sub population.

**S8 Fig.**
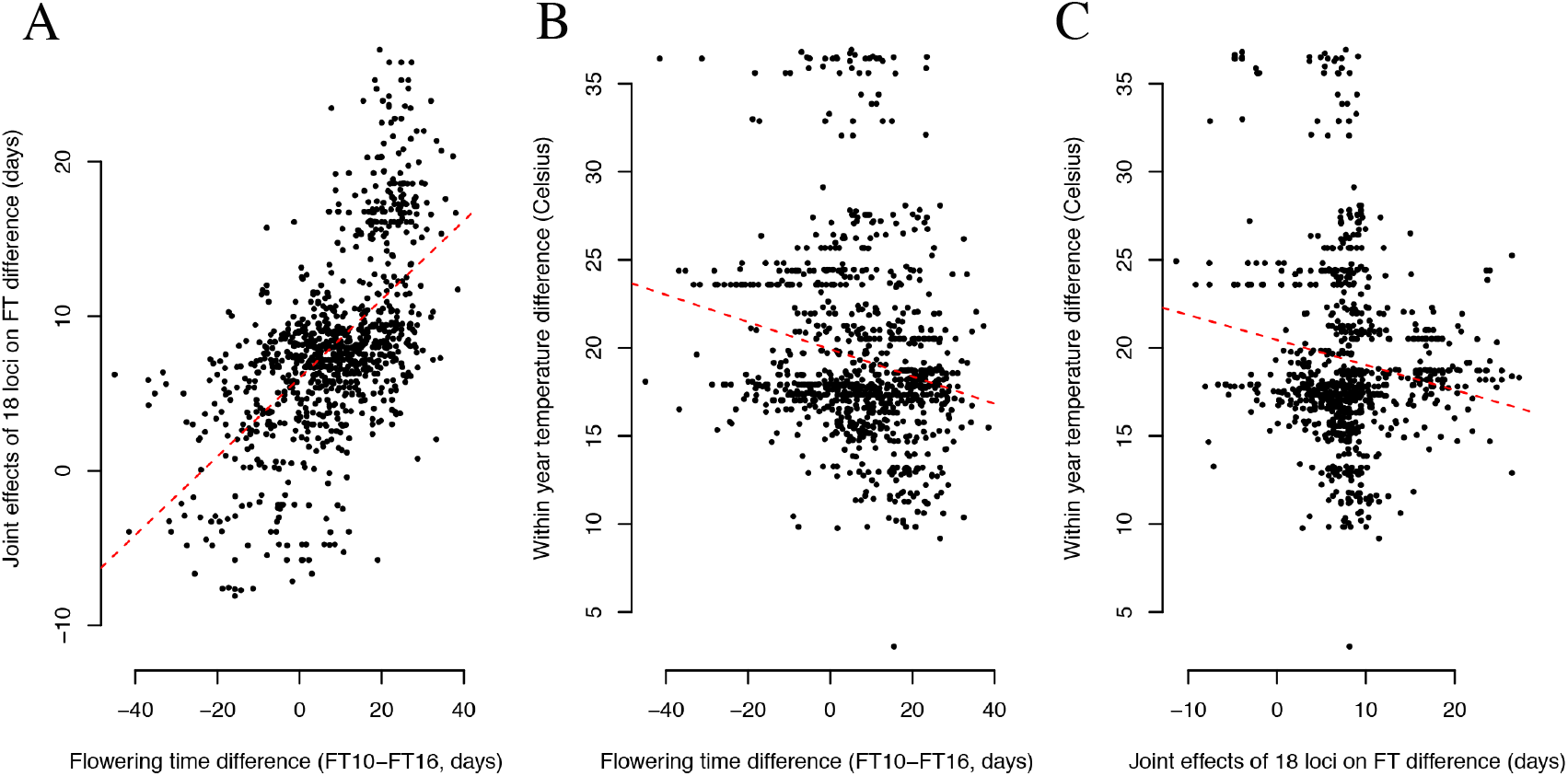
Correlations between observed and modelled flowering time differences for accessions grown at 10°C and 16°C and the within year temperature fluctuations at their sampling locations. **(A)** The correlation between the measured flowering time difference (FT10-FT16) and the flowering time difference modelled by the joint effects the 18 loci with significant genotype by temperature interactions (Adjusted R-square = 0.34; P-value = 1.3 × 10^−90^; Fig 2A). (**B**) There is a significant correlation (Adjusted R-squared = 0.04; P-value = 1.7 × 10^−10^) between the within year temperature fluctuation (temperature difference between hottest and coldest day) at the sampling locations of the accessions [22] and the measured flowering time difference (FT10-FT16). (**C**) There is also a significant correlation (Adjusted R-squared = 0.03; P-value = 1.2 × 10^−7^) between the within year temperature fluctuation (temperature difference between hottest and coldest day) [22] and the modelled flowering time differences for the accessions from the joint effects of the 18 loci with significant genotype by temperature interactions (Fig 2A).

